# Underlying mechanisms and ecological consequences of variation in exploratory behavior of the Argentine ant, *Linepithema humile*

**DOI:** 10.1101/369132

**Authors:** Hannah Page, Andrew Sweeney, Anna Pilko, Noa Pinter-Wollman

## Abstract

Uncovering how and why animals explore their environment is fundamental for understanding population dynamics, the spread of invasive species, species interactions etc. In social animals, individuals within a group can vary in their exploratory behavior and the behavioral composition of the group can determine its collective success. Workers of the invasive Argentine ant (*Linepithema humile)* exhibit individual variation in exploratory behavior, which affects the colony’s collective nest selection behavior. Here we examine the mechanisms underlying this behavioral variation in exploratory behavior and determine its implications for the ecology of this species. We first establish that individual variation in exploratory behavior is repeatable and consistent across situations. We then show a relationship between exploratory behavior and the expression of genes that have been previously linked with other behaviors in social insects. Specifically, we find a negative relationship between exploratory behavior and the expression of the FOR gene. Finally, we determine how colonies allocate exploratory individuals in natural conditions. We find that ants from inside the nest are the least exploratory individuals, while workers on newly formed foraging trails are the most exploratory individuals. Furthermore, workers in the spring, when new resources emerge, are more exploratory than workers in the winter, potentially allowing the colony to find and exploit new resources. These findings reveal the importance of individual variation in behavior for the ecology of social animals and provide mechanistic and functional explanations for how such behavioral variation can emerge.

## Introduction

Exploratory behavior is fundamental for how animals interact with their environment. Animals explore new environments when they disperse from their natal habitat (Stamps, 2001), expand their home range (Onen and Hanski, 2006), and search for new resources (Kramer and Weary, 1991). For example, exploratory individuals of invasive species can be responsible for expanding the invaded range (Cote et al., 2011; Fogarty et al., 2011). Despite the advantages of exploring for new resources, there are many associated costs, including spending valuable energy during the search process (Stamps et al., 2005), encountering novel predators (Eliassen et al., 2007; Kramer and Weary, 1991), and ending up in worse habitat than the one the animal departed from (Stamps, 2001).

Social animals can benefit from the discovery associated with exploration with minimal costs because individuals in social groups vary in their behavior. Thus, non-exploratory group members may benefit from new information obtained by risk-taking exploratory group members, without paying the costs associated with obtaining the new information. For example, in bird flocks, ‘scroungers’ access resources found by ‘producers’ (Aplin and Morand-Ferron, 2017; Giraldeau et al., 1994; Katsnelson et al., 2008). Individuals in a group could be competing over resources and so studies of producer-scrounger dynamics often focus on competition rather than the potential mutual benefits (Beauchamp and Giraldeau, 1996; Koops and Giraldeau, 1996). In social insects, selection acts at the level of the colony, and so if certain individuals can enhance the colony’s ability to obtain resources, the entire colony benefits. For example, in honey bees, certain foragers act as ‘scouts’ (Biesmeijer and de Vries, 2001) who are more likely than other foragers to locate new resources (Liang et al., 2012). When the scouts find new resources, they recruit other foragers to the location of these resources (von Frisch, 1967), thus improving overall colony foraging success, especially in patchy environments (Anderson, 2001; Beekman and Bin Lew, 2008; Dechaume-Moncharmont et al., 2005; Donaldson-Matasci and Dornhaus, 2012; Dornhaus and Chittka, 2004).

Animals vary consistently in their behavior (Sih et al., 2004) and these consistent behavioral differences have long been studied in social insects (Beshers and Fewell, 2001; Gordon, 1996; Jaisson et al., 1988; Jandt et al., 2014; Oster and Wilson, 1978). Work on social insects has focused primarily on differences among individuals in which tasks they perform, such as foraging, nursing, nest maintenance, etc. However, there is also variation in the way that each individual performs a task, for example, some individuals may be diligent in their performance of any task while others perform tasks only intermittently and spend much of their time resting (Charbonneau and Dornhaus, 2015; Pinter-Wollman et al., 2012; Robson and Traniello, 2002). This behavioral variation among individual workers can have great implications on the collective behavior and success of the colony (Hui and Pinter-Wollman, 2014; Pinter-Wollman, 2012; Pruitt and Riechert, 2011). Yet, only little is known about the mechanisms that underlie the behavioral differences among individuals and about the broad ecological implications of these behavioral difference (Beekman and Jordan, 2017).

One potential mechanism that can underlie variation in behavior is genetic variation, specifically, variation in the expression of particular genes, which allows for both flexibility and persistence (Bell and Aubin-Horth, 2010; LeBoeuf and Grozinger, 2014). Behavioral genomics has uncovered gene candidates that control social behavior (Robinson et al., 2005; Smith et al., 2008; Zayed and Robinson, 2012). For example, the honeybee model system has provided extraordinary insights into the genes whose expression regulates the division of labor between foraging (Ben-Shahar et al., 2002), scouting (Liang et al., 2012), nursing (Whitfield et al., 2003), and defense (Alaux and Robinson, 2007). One well studies gene is the FOR gene, which positively associates with foraging behavior in honeybees (Ben-Shahar et al., 2002) and negatively associates with foraging in harvester ants (Ingram et al., 2005).

The Argentine ant (*Linepithema humile)* has invaded ecosystems throughout the world with remarkable success (Suarez et al., 2001). Workers of *L. humile* vary in their exploratory behavior, and a group’s composition of exploratory behavior determines its ability to select a suitable nest site (Hui and Pinter-Wollman, 2014). Colonies of *L. humile* occupy more than one nest site (i.e., are polydomous) and expand the number of nest sites they occupy during the spring, when resources are abundant (Heller and Gordon, 2006). *L. humile* establish persistent foraging trails to long-lasting food sources and in between nests and they form new smaller trails to new ephemeral resources (Flanegan et al., 2013). Uncovering the mechanisms that underlie the collective exploration of *L. humile* can help predict where they will spread to next and aid mitigation of further spread.

Here we ask if individual variation in exploratory behavior is persistent and whether colonies of *L. humile* allocate exploratory individuals to where they are most needed and during the times of year when they are most beneficial to the colony. First, we examine if individual workers maintain the same exploratory behavior over days and across different assays for exploratory behavior. We then ask if there is a relationship between exploratory behavior and the expression of the FOR gene. We predict that because foraging behavior requires extensive movements outside the nest and the FOR gene is downregulated in foraging ants (Ingram et al., 2005), we will find a negative relationship between exploratory behavior and the expression of the FOR gene. Finally, we ask where and when can exploratory individuals be found in a natural setting. If colonies allocate exploratory individuals to where they are most needed, we expect to find the least exploratory individuals inside the nest and the most exploratory individuals on newly formed foraging trails. We further expect to find more exploratory individuals in the spring, when colonies expand their range, compared to the winter.

## Methods

### Exploratory behavior

Exploratory behavior of individual ants was quantified using an eight-arm maze with spices at the end of each arm, following the methods in (Hui and Pinter-Wollman, 2014; Modlmeier and Foitzik, 2011) (Figure 1A). An ant was placed in the center chamber of the maze for 5min and we recorded the total number of visits it made to any spice as its exploratory behavior. A visit was defined as an ant moving at least one body length into the maze arm leading to a spice. At the end of each trial we cleaned the apparatus with ethanol and removed and replaced spices that were touched by the ant.

**Figure 1:**
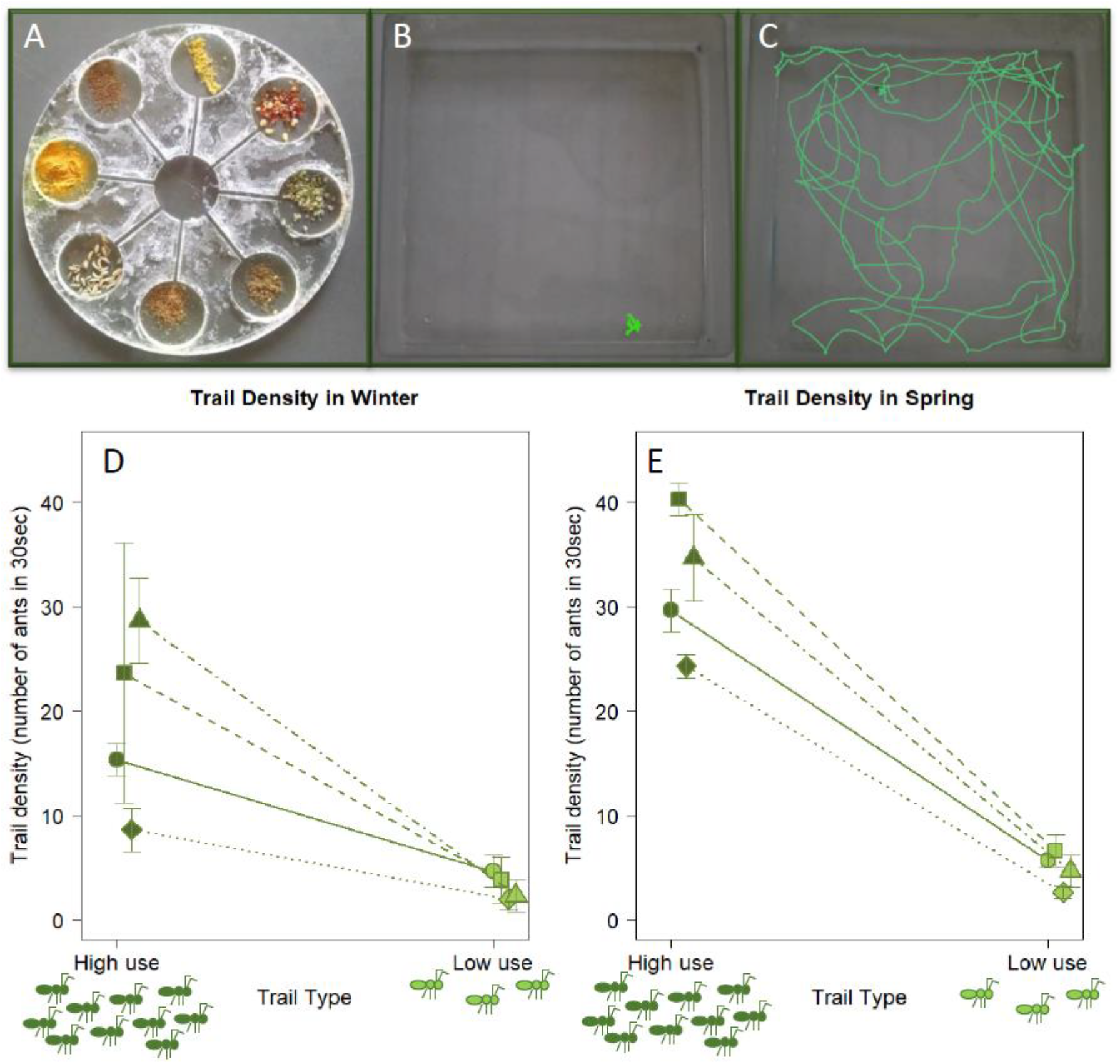
Quantifying exploration and trail use. (A) The 8-armed maze used for quantifying exploration. (B-C) Walking trajectories (green lines) of ants walking in an open field for 5min. These ants made 0 (B) and 5 (C) visits to spices when tested in the 8-armed maze the previous day. (D-E) Trail density, measured as number of ants crossing an invisible line on the trail during 30 seconds. Values are averages of three counts on each high- and low-use trails, in the winter (D) and spring (E) in four colonies. Points are averages of three counts, bars are the standard deviation of these counts, and lines connect measures from the same colony. Points and error bars are slightly jittered along the x axis to improve visibility.

To examine if exploratory behavior is a persistent trait we collected 66 *L. humile* ants from a colony on the UCSD campus in October 2014. We housed the ants in individual containers, and tested their exploratory behavior three times, on three consecutive days, once each day. We determined the repeatability of exploratory behavior using ICC (Bell et al., 2009).

To determine if the 8-armed maze assay reliably captures exploratory behavior we compared the exploratory scores from the maze with the behavior of ants in an open field (Figure 1B, C). We collected 90 more *L. humile* workers on February 2015 from a colony on the UCSD campus and housed them in individual containers. On the first day after collection, each ant was tested in the 8-armed maze. On the second day, each individual was placed in an open plastic box (11×11cm) with fluon coating its walls to prevent the ant from escaping. We recorded the movement of the ant with a video camera for 5min. We then used the tracking software AnTracks (https://sites.google.com/view/antracks) to determine the length of each ant’s walking path during these 5min and its path tortuosity, measured as the standard deviation of the turning angle (Adler and Gordon, 1992). We then examined the relationship between distance traveled or path tortuosity with the number of spices visited in the 8-armed maze using Pearson’s correlation.

### Gene expression

To determine if gene expression varies with exploratory behavior, we examined the expression of the FOR gene in 18 individuals collected in January 2015 from a colony on the UCSD campus. The exploratory behavior of each individual was measured three times using the 8-armed maze, on three consecutive days to obtain an average exploratory behavior. Immediately after testing for exploratory behavior on the third day, ants were flash frozen using liquid nitrogen and their heads separated from the bodies on dry ice and under a microscope with a pair of pre-chilled forceps.

To extract RNA for qPCR analysis, individual heads were placed in a 1.5 ml microcentrifuge tube and homogenized with a glass pestle in 0.5 ml of TRIzol^®^ Reagent (Thermofisher Scientific Cat.No 15596-018). Total RNA was extracted using Zymo Research Direct-zol RNA MiniPrep (Zymo Cat. No. R2053) followed by DNAseI treatment on column following manufacturer’s instructions. Extracted RNA was dissolved in RNase-free water and stored at −80°C. The RNA concentration and purity of each sample (A260/280 ~1.9) was measured using a NanoDrop ND-1000 (ThermoFisher, USA). All RNA was assessed for integrity using a BioAnalyzer tapestation (Agilent Technologies), and samples with RNA Integrity Number (RIN) > 8.0 were used. 20 ng of total RNA were transcribed with qScript™ XLT cDNA SuperMix (Quanta Bioscience). cDNA was stored at −80 degrees and briefly transported on dry ice before being thawed and used for qPCR assay.

Primer sequences were designed to span introns of predicted *L. humile* inositol monophosphatase 2-like (LOC105675875, XM_012373351.1) mRNA allowing only the specific amplification of cDNA (F: CGGCAGCTCTTTACCAGTCG, R: GCGACTGGGATCTCGTCACT). A species specific 110bp fragment of glyceraldehyde-3-phosphate dehydrogenase 1 (LOC105675989, XM_012373525.1) was used as a reference gene (F: CGATTCCATGGGCAAAAGCC, R: AATGACTTTCTTCGCACCGC).

RT-qPCR with 2ng input cDNA per reaction in triplicates was done with BioRad CFX384 Real Time system in 10ul reaction volume using PerfeCTa^®^ SYBR^®^ Green FastMix^®^ Cat# 95072-250 followed by melting curve analysis to ensure homogeneity of the reaction product. RT-qPCR data was analyzed using BioRad CFX manager software and quantification of relative mRNA levels was calculated using the 2^-ΔΔCt method (Livak and Schmittgen, 2001).

Expression of the FOR gene was normalized by dividing its expression with that of the *L. humile* specific housekeeping gene ‘RNA polymerase II elongation factor Ell’, which exhibited low variability across samples (Thellin et al., 2009). We examined the relationship between exploratory behavior and the normalized expression of the FOR gene using Pearson’s correlation.

### Allocation of exploratory individuals in nature

To examine where exploratory individuals are found in natural colonies we collected *L. humile* workers from four colonies on the UCLA campus in 2017 (exact dates below) at three different locations per colony: (1) within the nest, (2) from low-use trails, and (3) from high-use trails. Trail usage was determined by the density of ants on the trails approximately ten feet from the nest entrance. To determine ant density we counted the number of ants that passed one point on the trail for 30 seconds. Each trail’s density was measured three times, once every 5min (Figure 1D, E). Trail usage was compared with a one sided, paired Wilcoxon signed-rank test for each season (see season definitions below). The four colonies sampled were at least 100m apart to ensure independent samples from functionally distinct colonies (Heller et al., 2008).

Approximately 30 individuals were collected first from the high-use trail, then from the low-use trail (approximately 10m from the nest entrance on each type of trail) and finally from inside the nest (90 ants total per colony). This order of collection was used to avoid any disturbance that the collection of ants from the nest might have had on trail usage. Ants were brought back to the lab the same day they were collected and each individual was stored in an individually-labelled cup with water and sugar water *ad lib* to ensure individual identification in repeated testing. The exploratory behavior of each ant was examined using the 8-armed maze twice over two days - once on each day. Because some ants died between the two days, at least 15 individuals were tested twice from each location. After the second trial was completed, all ants were released back to their nest.

To test for seasonal differences in exploratory behavior and in the allocation of exploratory individuals to various locations, we conducted the above procedure twice: first in the winter (February 3rd-March 10th, 2017) and then in the spring (April 18th-May 18th, 2017). The average temperature in the winter (14C), was three degrees C colder than in the spring (17C), and the air was drier in the winter (dew point 8.24C) compared with the spring (dew point 10.08C). Weather data were obtained from historical records from the closest weather station to UCLA, at Santa Monica (KSMO), through ‘Weather Underground’ (https://www.wunderground.com/history). Ants from the four colonies were collected in the same order during both seasons to ensure that the number of days between each seasonal collection was approximately equal for each colony.

Differences in exploratory behavior between seasons and collection locations were examined using a Generalized Linear Mixed Model (GLMM) with a log-link function using the lme4 R package (Bates et al., 2015). The dependent variable was the number of visits ants made to spices in the 8-armed maze during 5min. The fixed effects were collection location (nest, low-use, and high-use trails) and season (winter and spring) and the random effects were colony identity and the day on which the exploratory behavior of an ant was tested (first or second - each ant was tested twice). To determine the confidence of our estimates, we ran a Wald Chi-squared test using the Anova R function (‘MASS’ package (Venables et al., 2002)) on the GLMM results. Post-hoc comparisons between collection sites for each season were conducted with Tukey tests using the ‘lsmeans’ R package (Lenth, 2016). All analysis was conducted in R version 3.1.2 (R Core, 2014).

## Results

### Exploratory behavior

Exploratory behavior is a repeatable trait. The 66 ants whose exploratory behavior was examined over three consecutive days had a repeatability score of 0.28 (CI: 0.13, 0.44) for the number of visits that ants made to spices, measured with ICC. This repeatability score is comparable to that found in other animals (Bell et al., 2009).

Exploratory behavior quantified with the 8-armed maze reflected the ants’ movement in an open field. Distance moved significantly and positively correlated with the number of visits an ant made to spices in the 8-armed maze in the preceding day (Pearson’s correlation: r = 0.28, p = 0.01; Figure 2). Interestingly, we did not detect a significant relationship between an ant’s path length and its tortuosity (Pearson’s correlation: r = −0.16, p = 0.16), nor was there a significant relationship between path tortuosity and the number of visits to spices in the 8-armed maze (Pearson’s correlation: r = 0.03, p = 0.77).

**Figure 2:**
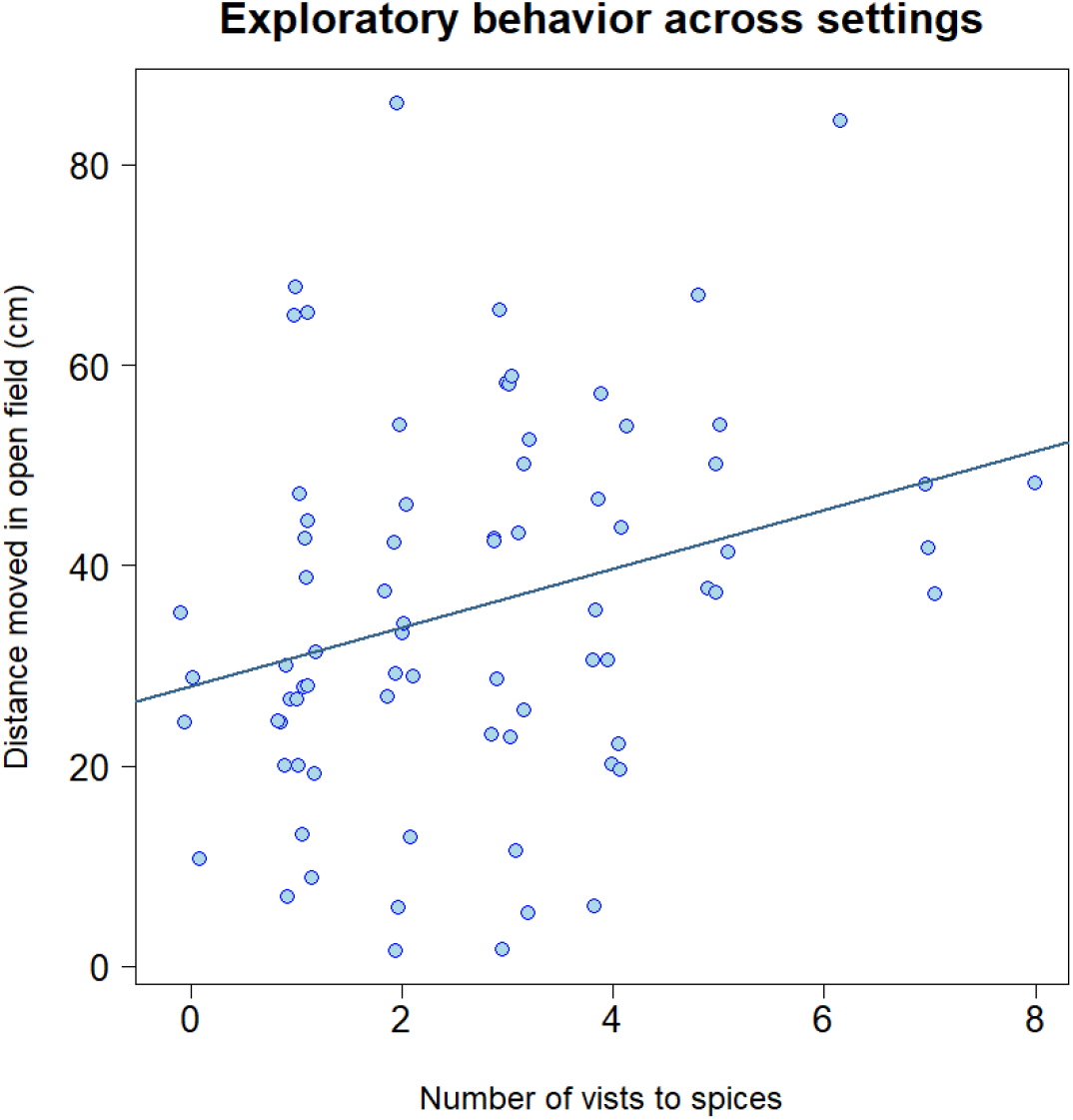
Exploratory behavior across settings. A significant positive correlation between the exploratory behavior of an ant when measured in the 8-armed maze and the total distance it traveled in an open field. Points are slightly jittered along the x axis to improve visibility.

### Gene expression underlies exploratory behavior

Exploratory behavior is related to the expression of the FOR gene. We found a significant negative correlation between the number of visits an ant made to spices in the 8-armed maze and the expression of the FOR gene (Pearson’s correlation r = −0.59, p = 0.01; Figure 3).

**Figure 3:**
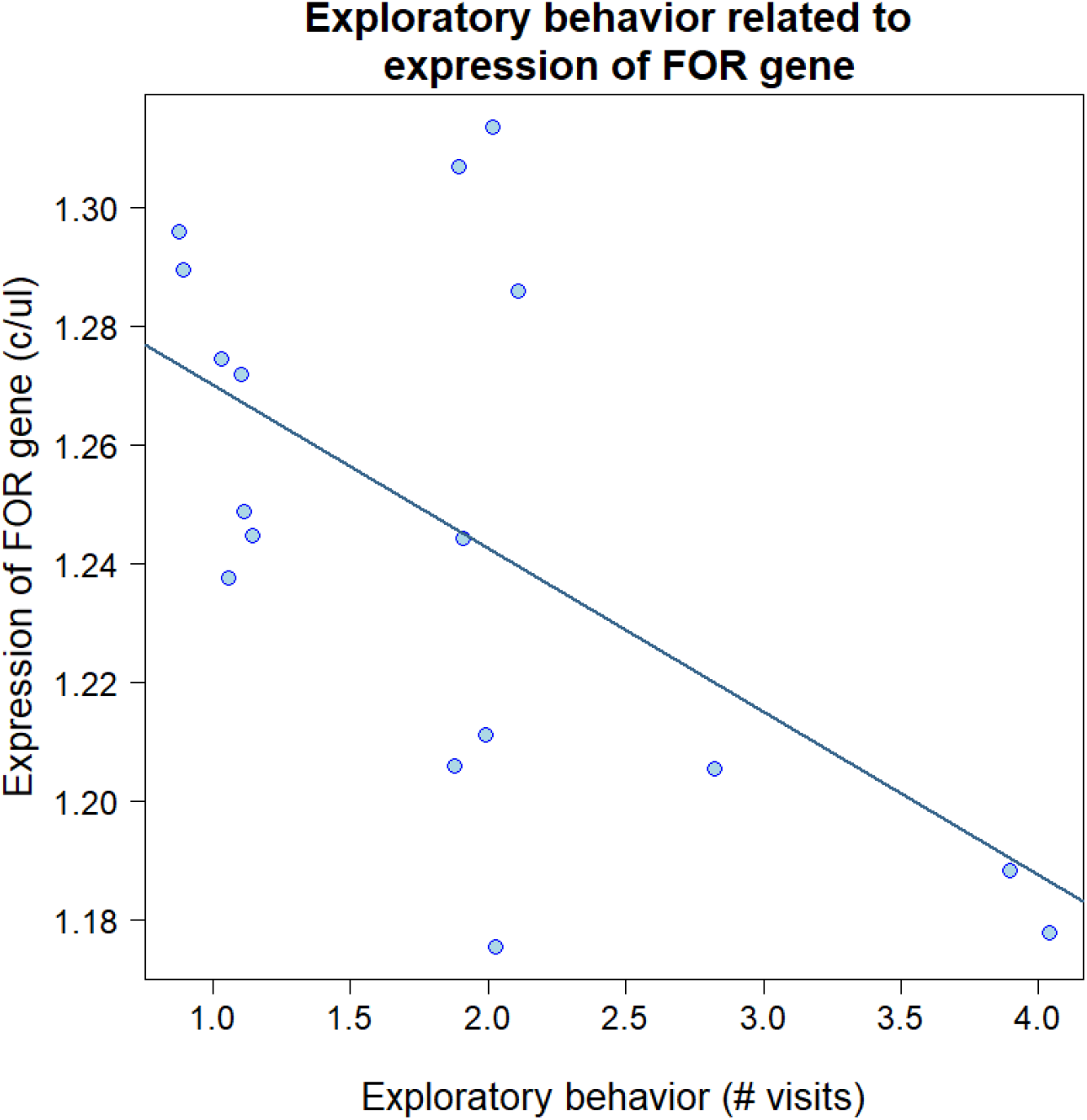
Exploratory behavior related to gene expression. A significant negative correlation between exploratory behavior (number of visits to spices in the 8-armed maze) and the expression of the FOR gene (copy number per ul, corrected for expression of housekeeping genes). Points are slightly jittered along the x axis to improve visibility.

### Allocation of exploratory individuals in nature

*L. humile* colonies allocate exploratory individuals to where they are most needed. High-use foraging trails were significantly denser than low-use trails in both the winter and spring (paired, one sided Wilcoxon test: winter: V = 120, p = 0.0003; spring: V = 78, p = 0.001; Figure 1D, E). In both seasons, the most exploratory individuals were found on the low-use trails and the least exploratory individuals were found inside the nest. Individuals on the high-use foraging trail exhibited intermediate levels of exploratory behavior in the winter and exploratory behavior similar to workers from inside the nest in the spring (GLMM location effect: χ^2^= 53.64, DF = 2, p < 0.0001; Figure 4). Seasonally, ants were significantly more exploratory in the spring than in the winter (GLMM season effect: χ^2^ = 12.43, DF = 1, p < 0.0005; Figure 4). The random effects had very little impact on the model (variance for ‘colony’ = 0.01, and for ‘day on which the ant was tested’ = 0.02).

**Figure 4:**
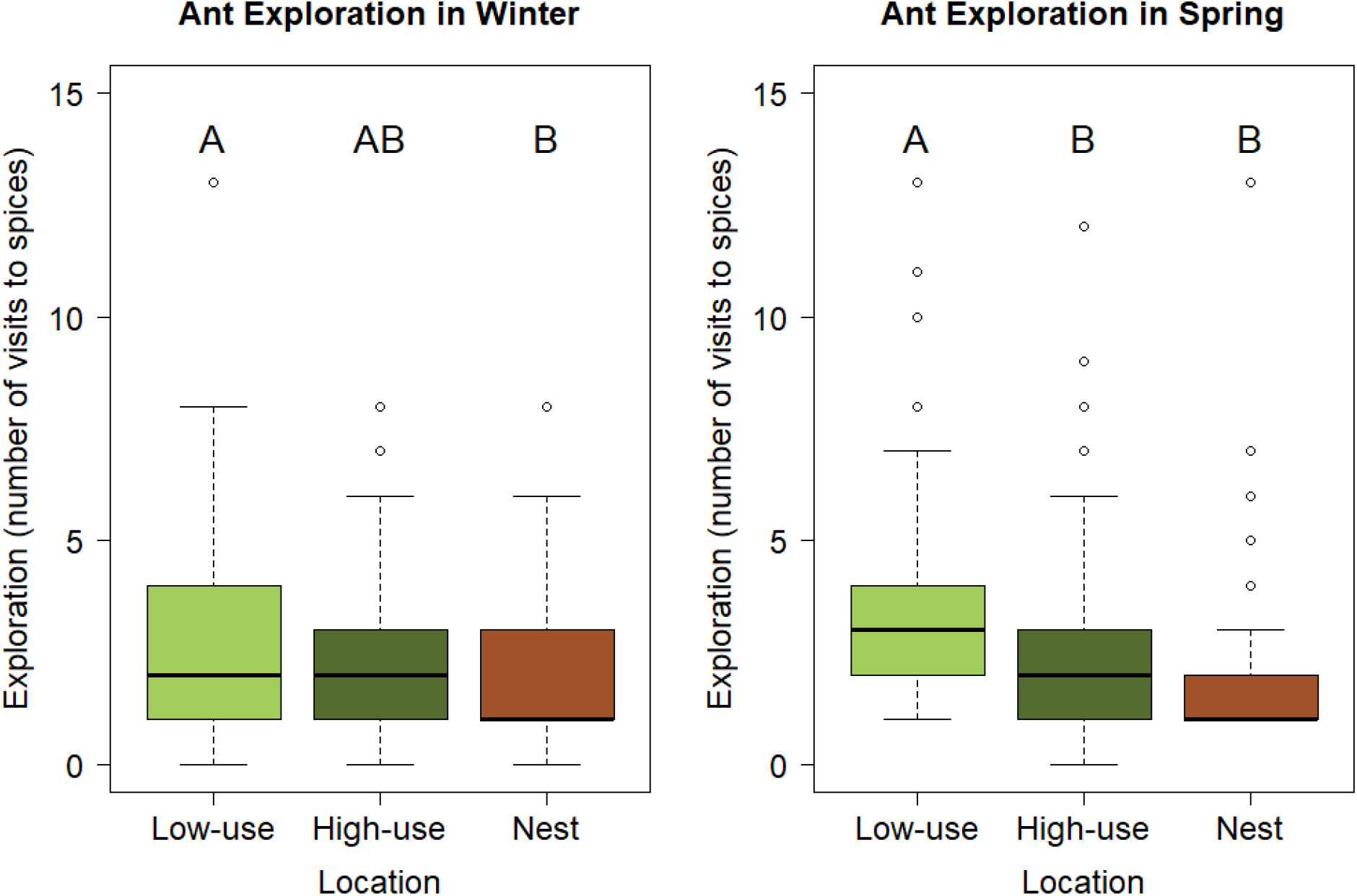
Exploratory behavior in nature. Exploratory behavior of *L. humile* workers collected from different locations: low- and high-use foraging trails, and from inside the nest in the winter (left) and the spring (right). Boxes indicate the lower and upper quartiles; horizontal lines within boxes indicate the median; whiskers extend to the 1.5 interquartile range from the box; and open circles indicate outliers. Boxes that do not share a letter are statistically different according to a post-hoc Tukey test (p-value < 0.01).

## Discussion

The amount of exploration an *L. humile* worker exhibits is a repeatable trait that is mediated, at least partially, by the expression of the FOR gene. In addition to identifying the underlying mechanisms that determine exploratory behavior of individual workers, we showed that colonies allocate exploratory individuals to where they are most needed ecologically. Thus, our work links individual and collective behaviors mechanistically and functionally from the level of the genes through individuals to the society.

Many behavioral traits are repeatable, however, the mechanisms that underlie such repeatability are understudied. The exploratory behavior of Argentine ants was found to be as repeatable as some behaviors studied in other animals (Bell et al., 2009). Thus, on a time scale of a few days, individuals maintain a certain level of exploratory behavior. Further work is required to determine if exploration changes on longer time scales, for example, as individuals age or when they switch their task, e.g., from nurse to forager. We identified that the expression of the FOR gene decreases with exploratory behavior. The effect of gene expression on a behavior allows for behavioral persistence on certain timescales and flexibility on other time scales. For example, exploratory behavior may persist as long as the appropriate level of expression of the FOR gene is maintained, and when this expression changes, the behavior may change as well. Foragers of harvester ants have the lowest expression of the FOR gene compared to workers performing other tasks (Ingram et al., 2005). Foragers spend more time than any other workers outside the nest because they travel to retrieve food. Thus, the low expression of the FOR gene in exploratory *L. humile*, who travel longer distance than other workers, is consistent with its low expression found in foragers. Interestingly, an opposite trend is found in the red imported fire ant, *Solenopsis invicta.* In that system, the FOR gene is over-expressed in foragers and colonies with highly exploratory individuals express more of the FOR gene (Bockoven et al., 2017). Thus, species differ in the way that gene expression relates to behavior. A recent study (Ingram et al., 2016) found both daily oscillations in the expression of the foraging gene as well as differences among tasks. Changes that occur in gene expression at the temporal scale of hours are influenced by the amount of light that foragers (but not nurses) are exposed to (Ingram et al., 2016). Thus, it is possible that the exposure of exploratory individuals to light when they are outside the nest may control the expression of the FOR gene. This hypothesis may be tested by comparing the FOR gene expression between exploratory individuals from low-use trails and less exploratory individuals from high-use trails. Both of these workers potentially have a similar exposure to light but they differ in their exploration.

Exploratory behavior has ecological significance at both the individual and collective levels. First, we showed that exploratory behavior, quantified in a simple 8-armed maze, is equivalent to the, more ecologically relevant actual distance that an ant travels in an open space (Figure 2). Interestingly, path tortuosity did not relate to distance traveled in an open field or to the exploration of an 8-armed maze. Previous work showed that as the path tortuosity of ants’ paths increases, interaction rate decreases (Pinter-Wollman et al., 2011). Thus, our inability to detect a relationship between path tortuosity and exploratory behavior suggests that exploratory behavior does not necessarily affect interaction patterns, which are known to regulate collective actions, such as foraging activity (Greene and Gordon, 2007; Pinter-Wollman et al., 2013). Thus, the amount of exploratory behavior exhibited by ants in a colony, likely has a greater impact on the area a colony can reach, because exploratory ants travel far, rather than on how the colony coordinates its activities.

Exploratory individuals were allocated to where they were most needed and at the appropriate time of year. In natural conditions, we expect ants that have high exploratory behavior to travel to remote locations and cover more ground than other individuals. Therefore, we expected that exploratory individuals will be allocated to where long-distance traveling is most needed. Indeed, we found that ants on low-use trails were more exploratory than those on high-use trails. Half of the low-use trails went into trees (personal observations), where ants often feed on honeydew from aphids. Aphids can move around, and so it is possible that exploratory behavior would help locate them. In contrast, high-use trails went towards trash-cans (personal observations), which are a reliable, constant, food source that does not move, and therefore does not require much exploration. Previous work on high-use trails (Flanegan et al., 2013) showed that certain individuals will occasionally leave these trails and find new resources that are adjacent to the trail. It is possible that these meandering ants are more exploratory than individuals on the main trail and that variation in exploratory behavior on high-use trails is beneficial for finding new resources along the path to an established resource.

Tasks performed inside the nest, such as nursing and cleaning, likely do not require as much exploratory behavior as tasks performed outside the nest, such as seeking for new food sources, because the space inside the nest is smaller than outside. Indeed, ants outside the nest, especially on low-use trails, were significantly more exploratory than ants inside the nest. It is possible that the similarity between exploratory behavior on high-use trails and inside the nest, especially in spring (Figure 4) is a result of the way we sampled ants from inside the nest. If many foragers are waiting near the nest entrance to be recruited, sampling ants from the top portion of the nest, as we did here, would potentially include a greater proportion of foragers than samples from deeper in the nest that may reach more brood caretakers. Thus, sampling ants from deeper inside the nest might have produced a greater difference in exploratory behavior between ants collected in the nest and on high-use trails, however, such sampling would be destructive and might have disturbed colony activity.

Finally, exploratory behavior was more predominant in the spring, when more food is available and when *L. humile* colonies expand their local range (Heller and Gordon, 2006), compared to the winter. Thus, colonies allocate exploratory individuals both spatially and temporally, based on the collective ‘needs’ of the colony. It is possible that when the weather is warmer ants move faster and therefore are more exploratory. It is further possible that temperature plays a role in the expression of the FOR gene, just as exposure to light does (Ingram et al., 2016). Further studies of the proximate mechanisms that underlie differences in behavior at the individual will uncover the ways in which the emergent collective behavior of the colony responds to changes in its environment.

By linking individual and collective behaviors, using both proximate and ultimate perspectives, our work brings us closer to uncovering how collective behaviors emerge. The genetic, mechanistic, control of variation in behavior among individuals acts in conjunction with the functional, adaptive, requirements of the colony for workers that express different behaviors in different locations. Further work on the feedback between how environmental pressures affect the genetic mechanisms and how the genetic mechanisms affect behavioral responses to environmental changes are necessary to elucidate the evolution of behavior.

## Author contributions

HP, AS, and NPW designed the study. HP and AS collected the behavioral data, AP conducted molecular work. HP and NPW analyzed the data and wrote the first draft of the paper.

## Data Availability

Data will be made available on Dryad upon acceptance of the paper.

